# BRD4 isoforms have distinct roles in tumor progression and metastasis in embryonal rhabdomyosarcoma

**DOI:** 10.1101/2023.07.26.550665

**Authors:** Dipanwita Das, Jia Yu Leung, Vinay Tergaonkar, Amos Hong Pheng Loh, Cheng-Ming Chiang, Reshma Taneja

## Abstract

BRD4, a bromodomain and extraterminal (BET) protein, is deregulated in multiple cancers and has emerged as a promising drug target. However, the function of the two main BRD4 isoforms (BRD4-L and BRD4-S) has not been analyzed in parallel in most cancers. This complicates determining therapeutic efficacy of pan-BET inhibitors. In this study, using functional and transcriptomic analysis, we show that BRD-L and BRD4-S isoforms play distinct roles in embryonal rhabdomyosarcoma. BRD4-L has an oncogenic role and inhibits myogenic differentiation, at least in part, by activating myostatin expression. Depletion of BRD4-L *in vivo* impairs tumor progression but does not impact metastasis. On the other hand, depletion of BRD4-S has no significant impact on tumor growth, but strikingly promotes metastasis *in vivo*. Interestingly, BRD4-S loss results in the enrichment of BRD4-L and RNA Polymerase II at integrin gene promoters resulting in their activation. Our work unveils isoform-specific functions of BRD4 and demonstrates that BRD4-S functions as a gatekeeper to constrain the full oncogenic potential of BRD4-L.

## Introduction

Rhabdomyosarcoma (RMS), one of the most common soft tissue sarcomas among pediatric patients, arises due to a block in myogenic differentiation. RMS tumour cells fail to differentiate terminally despite expression of core myogenic transcription factors (1). RMS is traditionally classified into two major subtypes: alveolar (ARMS) and embryonal (ERMS), which account for 20% and 70% of RMS cases respectively (2,3). PAX3-FOXO1 and PAX7-FOXO1 fusion proteins are present in about 70%-80% of ARMS but absent in ERMS. ARMS with fusion positive (FP) status is associated with worse prognosis. Patients with fusion negative ARMS have clinical outcomes similar to ERMS (4,5).

In ERMS, there is a loss of heterozygosity on the short arm of chromosome 11(11p15.5), leading to inactivation of tumour suppressor genes (6). The overall survival rate for patients with relapsed or metastatic RMS remains as low as 21% and 30% respectively. RMS is characterized by an aberrant epigenetic landscape. Altered expression of DNA methyltransferases and demethylases, microRNAs and enzymes involved in histone methylation, phosphorylation and acetylation has been observed. This leads to altered expression of genes involved in cellular proliferation, DNA replication, differentiation, epithelial-mesenchymal transition (EMT) and tumour progression. Epigenetic reprogramming thus provides an opportunity to identify novel druggable targets (7).

Bromodomain-containing protein 4 (BRD4), a member of bromodomain and extraterminal (BET) family, is an epigenetic regulator that plays an important role in embryogenesis and cancer development (8). The BET family proteins are acetyl-lysine readers, which primarily bind to acetylated chromatin and transcription factors (9). The BET proteins are characterized by two tandem bromodomains (BD1 and BD2), which bind acetylated lysine residues on target proteins (10). BRD4, along with other BET proteins, accumulates on active transcription regulatory elements and enhances gene transcription in both the initiation and elongation phase (11).

BRD4 gene encodes two major naturally occurring spice variants: BRD4-long (BRD4-L) and BRD4-short (BRD4-S) isoforms. The two variants are generated in a constant and balanced ratio to ensure homeostatic functioning of the protein (12). BRD4-L has an extended proline-rich region and a positive transcription elongation factor (P-TEFb)-interacting region at its C-terminal motif (CTM; see Fig. 1A), while BRD4-S with phase-separation properties, functions to organize chromatin and transcription factors for activation of gene transcription (13). BRD4-L and Mediator together form the transcription initiation complex, which along with P-TEFb then phosphorylates dual serine residues (Ser5 and Ser2) in RNA Polymerase (Pol) II and modulates transcription initiation and early elongation (11,14). BRD4-S, on the other hand, contains three unique C-terminal residues, GPA, derived from an alternatively spliced C-terminal exon (9).

**Figure 1:**
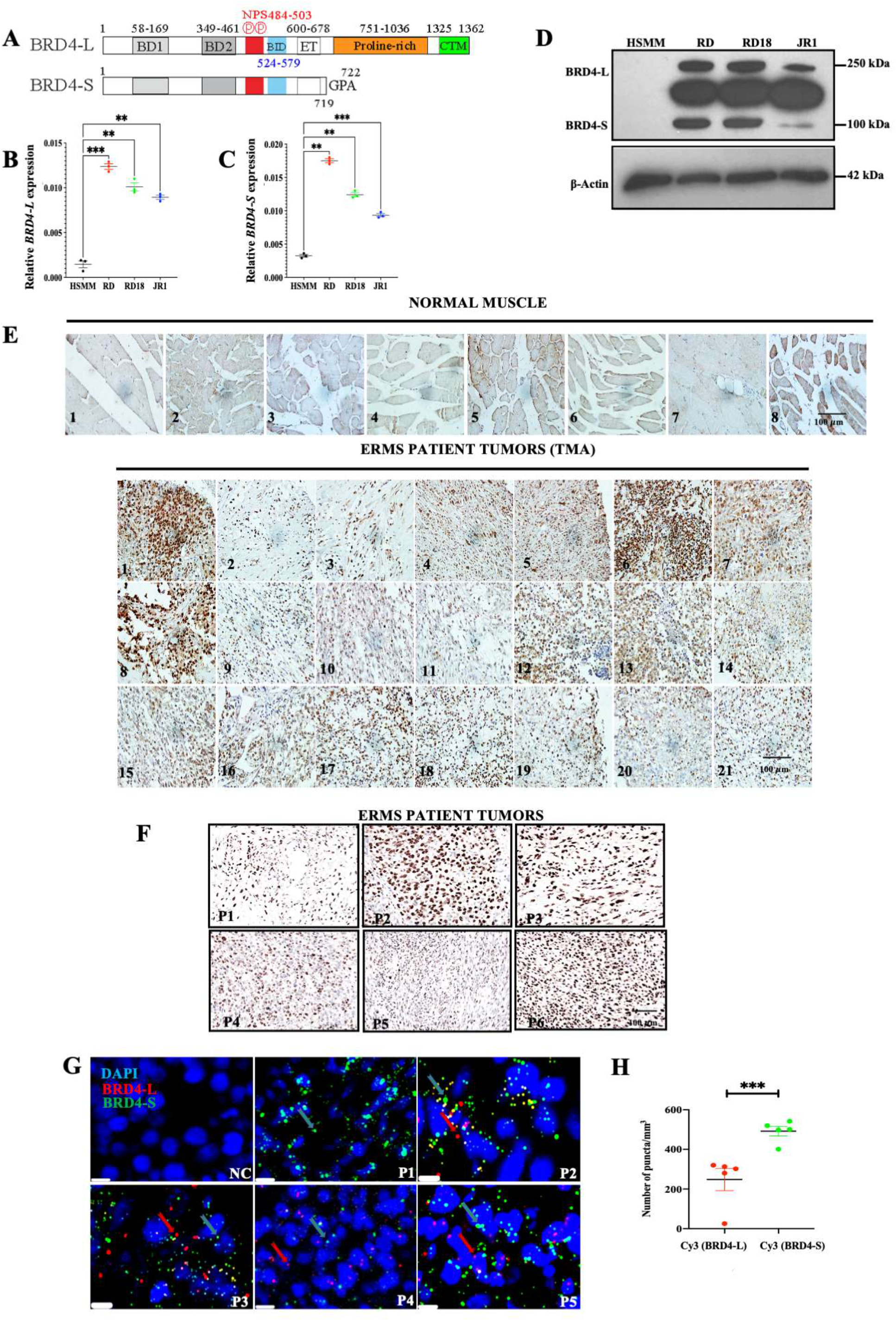
BRD4 is overexpressed in ERMS. **(A)** Schematic presentation of BRD4-L and BRD4-S domain structure. **(B-C)** BRD4-L and BRD4-S mRNA levels were examined in HSMM, RD, RD18 and JR1 by RT-qPCR analysis. Values correspond to the average ± SEM (n=3). Statistical significance was calculated by one-way ANOVA analysis. **p ≤ 0.01, ***p ≤ 0.001. **(D)** Western blot analysis showing expression of BRD4-L and BRD4-S isoforms in HSMM, RD, RD18 and JR1 cell lines. β-Actin was used as loading control. Images are representative of at least three biological replicates. **(E)** TMA consisting of 8 normal skeletal muscle samples and 21 ERMS patient tumors was analyzed by IHC using anti-BRD4 antibody. Images were taken at 40X magnification. Scale bar: 100μm. **(F)** 6 archival ERMS tumor specimens (P1-P6) were analysed by IHC using anti-BRD4 antibody. Images were taken at ×40 magnification. Scale bar: 100μm. **(G)** Detection of BRD4-L and BRD4-S RNA transcripts in ERMS patient tumor specimens (P1-P5) using RNAscope assay. At least 3 fields were imaged for each patient sample and one representative image is shown. Nuclei were stained with DAPI (blue). Scale bar: 20 µm. **(H)** BRD4-L and BRD4-S transcripts were quantified as red and green puncta respectively and represented as a scatter plot. A field with approximately 50 nuclei was chosen for quantification. No signal was observed in the negative control (NC) slide. Values correspond to the average ± SEM. Two-tailed non-parametric unpaired t test was performed for statistical analysis. ***p≤0.001.

BET inhibitors (BETi) are in pre-clinical and clinical trials for the treatment of various cancers (15). Targeted inhibition of BRD4 suppresses tumour growth in breast and prostate cancer, as well as in acute myeloid leukemia and diffuse large B-cell lymphoma (16–19). Despite its success, BETi therapy is associated with acquired resistance upon prolonged treatment with the inhibitors (20). In addition, our recent studies have highlighted that BRD4-S and BRD4-L function in an opposing manner in triple-negative breast cancer (TNBC) by differential enhancer regulation of many gene networks, including the matrisome extracellular matrix (ECM) network (12). This highlights the importance of developing more target-selective BET inhibitors in cancer therapy.

Previous studies have identified the involvement of BRD4 in RMS. In FP-RMS, BETi JQ1 was found to disrupt PAX3-FOXO1 and BRD4 interaction, leading to degradation of the fusion protein (21). JQ1 reduces RMS tumour growth through anti-angiogenic mechanisms (22). The effect of JQ1 was directly proportional to the expression of MYC, regardless of whether the tumour was of embryonal or alveolar origin (23). In addition, a combination of BRD4 and PLK1 inhibitors showed synergistic anti-tumour effects in pediatric tumour models including RMS that is associated with MYCN-driven gene expression. Similarly, combined inhibition of BET proteins and metabolic tumour driver, mTORC1/2, was found to abrogate RMS growth (24). Despite growing evidence for a role of BRD4 in RMS, surprisingly, a functional distinction of the BRD4 isoforms in RMS remains elusive.

In this study, we examined the role of BRD4-L and BRD4-S in ERMS. We show that BRD4-L and BRD4-S isoforms are both expressed in ERMS cell lines and each isoform has a distinct role in tumor progression and metastasis. Transcriptomic and functional analysis *in vitro* and *in vivo* demonstrate that BRD4-L promotes proliferation and inhibits myogenic differentiation at least in part through regulation of myostatin. Interestingly, BRD4-S functions as a gatekeeper of BRD4-L in metastasis. When BRD4-S is depleted, metastasis is significantly enhanced and BRD4-L enrichment is apparent on several integrin gene promoters. Taken together, the results of this study highlight that BRD4 isoforms have specific functions in ERMS.

## Materials and Methods

### Cell lines, BRD4 transient and stable knockdown

ERMS cell lines (RD, RD18 and JR1) were a kind gift by Peter Houghton (Nationwide Children’s Hospital, OH, USA) and Rosella Rota (Bambino Gesu Children’s Hospital, ROM, IT). The cell lines were routinely tested for mycoplasma contamination using BioMycoX® Mycoplasma PCR Detection Kit (Atlantis Bioscience, SGP). RD cells were maintained in Dulbecco’s Modified Eagle’s Medium (DMEM) (Sigma-Aldrich, St. Louis, MO, USA), supplemented with 10% fetal bovine serum (FBS) (HyClone, Cytiva, USA) and 1% Penicillin-Streptomycin (HyClone, Cytiva, USA). RD18 and JR1 cells were cultured in RPMI 1640 with L-Glutamine (Thermo Fisher Scientific, Waltham, MA, USA) supplemented with 10% FBS and 1% Penicillin–Streptomycin. Primary human skeletal muscle myoblasts (HSMM) were purchased from Zen-Bio, Inc. (NC, USA) and cultured in skeletal muscle cell growth medium (#SKM-M, Zen-Bio, USA). HEK-293T cells were commercially purchased from American Type Culture Collection (ATCC) (Manassas, VA, USA) and cultured in DMEM supplemented with 10% FBS.

For transient knockdown, cells were transfected with 20 nM of siRNA using Lipofectamine RNAiMax (Thermo Fisher Scientific). Cells were analyzed 48 hr post transfection. Specific siRNAs for BRD4-L (targeting long variant; Cat. #: SI05044872), BRD4-S (targeting short variant 3’-UTR region; Cat#: SI05044865) and AllStars Negative control (Cat. #: SI03650318) were purchased from QIAGEN (MD, USA). ITGA4 siRNA (Cat. # SC 35685) and ITGA5 siRNA (Cat. # SC 29372) were purchased from Santa-Cruz Biotechnology Inc.

Stable knockdown cell lines were generated using lentiviral vectors. Around 90% confluent HEK-293T cells were transfected with packaging plasmid pIP1 (5 µg) and pIP2 (5 µg), envelope plasmid pIP/VSV-G (5 µg) (ViraPowerTM Lentiviral Packaging Mix, Thermo Fisher Scientific), 5 µg of lentiviral expression construct shRNA (pLKO.1 Mission shRNA DNA clone, Sigma-Aldrich Inc., St. Louis, MO, USA) control, shBRD4-L (#TRCN0000021424, Mission shRNA, Sigma-Aldrich Inc.) or shBRD4-S (#TRCN0000349782, Mission shRNA, Sigma-Aldrich Inc.) along with 30 µl of Lipofectamine 3000 (Thermo Fisher Scientific) following manufacturer’s instruction. The supernatant was replaced with Basal DMEM media 16 hr post-transfection. Viral pellet was collected and resuspended in DMEM media. RD cells were transduced with control lentiviral particles, shBRD4-L or shBRD4-S with polybrene (8 µg/ml) (Sigma-Aldrich Inc.). Transduced cells were preselected with 1 µg/ml puromycin (Sigma-Aldrich Inc.) for 4-5 days before expansion.

### RNAscope assay

RNAscope® Customized Probe-Hs-BRD4-O2 (BRD4-L; NM_058243.2; Target region 3314 - 4735) and Catalogue Probe-Hs-BRD4-O1 (BRD4-S; NM_014299.2; Target region 2381 - 3468) were obtained from Advanced Cell Diagnostics, Inc. (Hayward, CA) (Cat. #: 323100). Detection of BRD4-L and BRD4-S isoforms in formalin-fixed paraffin-embedded (FFPE) archival primary ERMS tumours was investigated using RNAscope® Multiplex Fluorescent v2 Assay following manufacturer’s instructions. A negative control (NC) slide provided in the kit was probed with a negative control probe targeting the DapB gene (Cat. #: 320871). Briefly, sections were deparaffinized, boiled with target retrieval reagents (30 min), digested with protease (40°C for 30 mins) and then hybridized with the probes (40°C for 2 hr). After six rounds of amplification, the probes were visualized with TSA Plus Cyanine3 (used for BRD4-L), and TSA Plus Cyanine5 (used for BRD4-S) (PerkinElmer, MA, USA. Coverslips were mounted with DAPI (Vectashield, Vector Laboratories, CA, USA) and images were acquired at 40X magnification with a Leica DCF 9000 GT digital camera using a Leica DMi8 microscope. For quantitative microscopical evaluation of control or test probe mRNA detection, QuPath software (v 0.4.1) was used to analyze each fluorophore channel separately and the absolute numbers of puncta were calculated in the separate channels.

### Proliferation, differentiation, migration and invasion assays

Proliferation was measured by using 5-bromo-2’-deoxy-uridine (BrdU) labelling and detection kit (Roche, BSL, CH) as described earlier (25). Cells were fixed and incubated with anti-BrdU antibody (1:100) after pulsing with 10 μM BrdU. Cells were incubated with anti-mouse Ig-fluorescein antibody (1:200) and mounted on glass slides using DAPI (Vectashield, Vector Laboratories, CA, USA). Images were acquired in 40X magnification using a fluorescence microscope BX53 (Olympus Corporation, Shinjuku, TYO, JP).

Transient or stable knockdown cells were cultured in differentiation media consisting of either basal DMEM (for RD cells) or RPMI 1640 (for RD18 and JR1 cells) with 2% Horse Serum (HyClone, Cytiva, USA) for 2–5 days as described (25). Cells were fixed with 4% paraformaldehyde, blocked and permeabilized with 10% horse serum and 0.1% Triton X containing PBS. Cells were then incubated with anti-MHC antibody (MHC; R&D Systems, Minneapolis, MN, USA) (1:400), followed by secondary goat anti-Mouse IgG (H + L) Highly Cross-Adsorbed Secondary Antibody, Alexa Fluor 568 (Thermo Fisher Scientific). Coverslips were mounted with DAPI (Vectashield, Vector Laboratories, CA, USA) and images were acquired at 40X magnification using a fluorescence microscope BX53.

Boyden chamber (Greiner Bio-One, KR, AT) assay was used to assess the migration and invasion as described (26). Cells were serum-deprived for 12 hr and 50 000 RD and RD18 cells or 30 000 JR1cells were seeded in serum-free media into the Transwell insert. After 24-30 hr, the inserts were stained with crystal violet and imaged at 10X magnification using a brightfield microscope (EVOS XL Core Imaging System, Thermo Fisher Scientific). Invasion was assessed using inserts coated with Matrigel (Corning, NY, USA) and seeded at a density of 70 000 cells/insert for RD and RD18; and 50 000 JR1 cells.

### RNA isolation, RT-qPCR and RNA-Sequencing (RNA-Seq)

Total RNA was extracted using Trizol agent (Thermo Fisher Scientific) and quantified by NanoPhotometer (Implen, CA, USA). 2 µg of total RNA was reverse-transcribed to single-stranded complementary DNA (cDNA) using iScript cDNA Synthesis Kit (Bio-Rad Hercules, CA, USA). RT-qPCR was run on Lightcycler 480 (Roche) using SYBR Green 1 Master Kit (Roche). PCR amplification was performed as described (26). CT values were normalized to the internal control GAPDH, and delta CT (ΔCT) values were obtained. Relative expression was calculated by 2^−ΔCT^ equation. RT-qPCR analyses were done in triplicate and each performed in three independent biological replicates. Primers for RT-qPCR are listed in Supplemental Table 1.

For RNA-Seq analysis, RD cells were transfected with siScr control, siBRD4-L or siBRD4-S siRNA for 48 hr. Total RNA was isolated and purified using RNeasy mini kit (QIAGEN). Quality of purified RNA was analysed using agarose gel electrophoresis and Agilent 2100. The samples were processed by Novogene AIT for complementary DNA (cDNA) library construction and read mapping. The raw image file from Illumina (HiSeq PE150) was transformed to Sequence Reads by CASAVA base recognition and stored in FASTQ (fq) format. Reads were filtered to gather clean reads using filtering conditions like reads without adaptors, reads containing number of bases that cannot be determined below 10% and at least 50% bases of the reads having Qscore denoting Quality value ≤5. For mapping of the reads, STAR software was used (27). 1 M base was used as the sliding window for distribution of the mapped reads. Differentially expressed genes (DEGs), Gene Ontology (GO) and Kyoto Encyclopedia of Genes and Genomes (KEGG) analysis was performed with corrected p value < 0.05 and fold change ≥ 1.2, as significant enrichment. RNA-Seq data has been deposited in GEO database under accession number GSE215393. The private token for reviewers is gtwnayygptkrjwr.

### Western blotting

Cells were lysed with RIPA lysis buffer (50 mM Tris HCl pH 7.5, 40 mM NaCl, 1% NP-40, 0.25% sodium deoxycholate, and 1 mM EDTA) containing protease inhibitor cocktail (Complete Mini, Sigma-Aldrich Inc.). Immunoblots were probed overnight with primary antibodies and incubated with relevant horseradish peroxidase (HRP)-conjugated secondary antibodies. The following primary antibodies were used: anti-BRD4 (short and long isoforms) (Cat. #: ab128874, Abcam (CB, UK), WB 1:1000), anti-BRD4 E2A7X (long isoform) (Cat. #: 13440S, CST (MA, USA), WB 1:1000), anti-GDF8/Myostatin (Cat. #: ab201954, Abcam, WB 1/1000), anti-MHC (Cat. #: sc-32732, Santa-Cruz Biotechnology Inc., WB 1:250), anti-MYOG (Cat. #: sc-12732, Santa-Cruz Biotechnology Inc., WB 1/250), anti-ITGA4 (Cat. #: 8440, CST, WB 1:1000), anti-ITGA5 (Cat. #: 4705, CST, WB 1:1000) and anti-β-actin (Cat. #: A2228, WB 1:10 000, Sigma-Aldrich Inc.). Appropriate secondary antibodies (IgG-Fc Specific-Peroxidase) of mouse or rabbit origin (Sigma-Aldrich Inc.) were used. Proteins were detected using Pierce ECL Western Blotting Substrate (Thermo Fisher Scientific).

### Chromatin immunoprecipitation (ChIP)

For standard ChIP-qPCR, 4×10^6^ RD, JR1, shBRD4-L, shBRD4-S or shScr control cells were fixed with formaldehyde as described (25). ChIP was conducted with 2 µg of IgG or purified antibodies against BRD4-L (Cat. #: 13440S, CST, 1:50), H3K9Ac (Cat. #: ab4441, Abcam) and RNA Pol II (Ser2P) (Cat. #: ab193468, Abcam) or 1 µg of BRD4-S (12) and analysed by qPCR using 4% of IP products and 0.2% of input DNA. Primers for ChIP-qPCR are listed in Supplemental Table 2.

### Reporter Assays

MyoD reporter activity was analysed as described (28). shScr, shBRD4-L and shBRD4-S cells were transfected with 200 ng of the MRF-dependent reporter 4Rtk-luc and 200 ng of MyoD in a 24-well plate format. 5 ng of Renilla reporter was co-transfected as an internal control. Transfection was carried out using Lipofectamine 3000 transfection reagent (Thermo Fisher Scientific). Reporter activity was analyzed 48 hr post-transfection with the Dual-Luciferase Reporter Assay System (Promega, Madison, WI, USA). Luminescence was analysed with a Varioskan plate reader using SkanIT software.

### Tissue Microarray and Immunohistochemistry

Tissue Microarray (TMA) (SO2082b) slides were purchased from US Biomax, Inc. (Derwood, MD, USA), which comprised of 27 ERMS tumour specimens and 8 striated muscle tissues. Six paraffin sections of archival primary ERMS tumours (P1-P6) from KK Women’s and Children Hospital in Singapore were also analyzed. Specimens were obtained following informed written consent under Institutional Review Board-approved protocol CIRB 2014/2079. Samples were analysed by IHC using anti-BRD4 antibody (Bethyl, MD, USA) (1:2000) using Dako REAL EnVIsion-HPR, Rabbit-Mouse kit (Dako, DK). Sections were counterstained with haematoxylin (Sigma-Aldrich Inc.). Slides were dehydrated and mounted using DPX (Sigma-Aldrich Inc.). For metastasis, paraffin sections of liver, lungs and kidney were stained with hematoxylin and eosin as described (26).

### Mouse Xenograft and Metastasis models

Animal procedures were approved by Institutional Animal Care and Use Committee under protocol number R20-1589. Six-week-old C.Cg/AnNTac-Foxn1nuNE9 female BALB/c nude mice (InVivos, Singapore) were injected subcutaneously in the right flank with one million (1×10^6^) of shScr, shBRD4-L or shBRD4-S RD cells (n=10/group). Tumour onset and growth were monitored and body weight was taken every alternate day. Tumour volume was calculated using the formula V = (L ×W×W)/2, where V is tumour volume; W is tumour width; and L is tumour length. Once tumours in the control group reached a size of 1.5 cm in diameter, mice were sacrificed and the resected tumours were used to prepare lysates for Western blot analysis.

To analyse metastasis, shScr, shBRD4-L or shBRD4-S RD cells (1 million cells/mouse) were injected in NOD/SCID mice (n=10/group) by tail vein injections. Body weights were taken every alternate day. The mice were sacrificed eight weeks after the injections. Resected tumours from the organs were used to prepare tumour lysates for Western blot analysis and the kidneys, lungs and liver were fixed in 4% PFA and processed for hematoxylin and eosin staining.

### Statistical Analysis

Biological assays are presented as mean ± standard error of the mean (S.E.M.). For statistical analysis, two-tailed unpaired Student’s t test for two-group comparison and one-way ANOVA with Tukey or Dunnett for multiple-group comparison using GraphPad Prism 9 (GraphPad software). P values < 0.05 were considered significant. Significance in all figures is indicated as follows: * p < 0.01, ** p < 0.05, ***p < 0.001, ****p < 0.0001 and n.s. is no significance. Each experiment was performed at least thrice as independent biological replicates.

## Results

### BRD4 is overexpressed in ERMS

Expression of BRD4-long (BRD4-L) and BRD4-short (BRD4-S) isoforms (**Fig. 1A**) was analyzed in patient-derived ERMS cell lines – RD, RD18 and JR1. An increased expression of both BRD4 isoforms in all the three cell lines was seen compared to primary human skeletal muscle myoblasts (HSMM) at both the mRNA and protein levels (**Fig. 1B-D**). Immunohistochemical analysis with an anti-BRD4 antibody that detects both isoforms showed that BRD4 expression was elevated in 21 ERMS patient tumours compared to 8 normal muscles in a tissue microarray (TMA) (**Fig. 1E**). Similarly, BRD4 expression was elevated in 6 archival ERMS tumour sections (**Fig. 1F**). To analyze the relative expression of BRD4-L and BRD4-S RNA transcripts in these ERMS tumor sections, we used RNAscope analysis. Quantification of the number of puncta for each isoform revealed that BRD4-S transcripts were more abundant compared to BRD4-L transcripts **(Fig. 1G, H)**.

### BRD4-L and BRD4-S isoforms have distinct roles in tumor progression and metastasis

RMS cells were recently reported to be sensitive to pan-BRD4 inhibitor JQ1 (23). We tested the impact of JQ1 treatment in RD cells. We observed a significant decrease in proliferation **(Fig. 2A, B)** and an increase in myogenic differentiation upon JQ1 treatment compared to DMSO-treated control cells **(Fig. 2C, D, E)**. However, unexpectedly invasion and metastasis were significantly increased with JQ1 treatment **(Fig. 2F, G, H),** suggesting that pan-BRD4 inhibition may have an undesired outcome.

**Figure 2:**
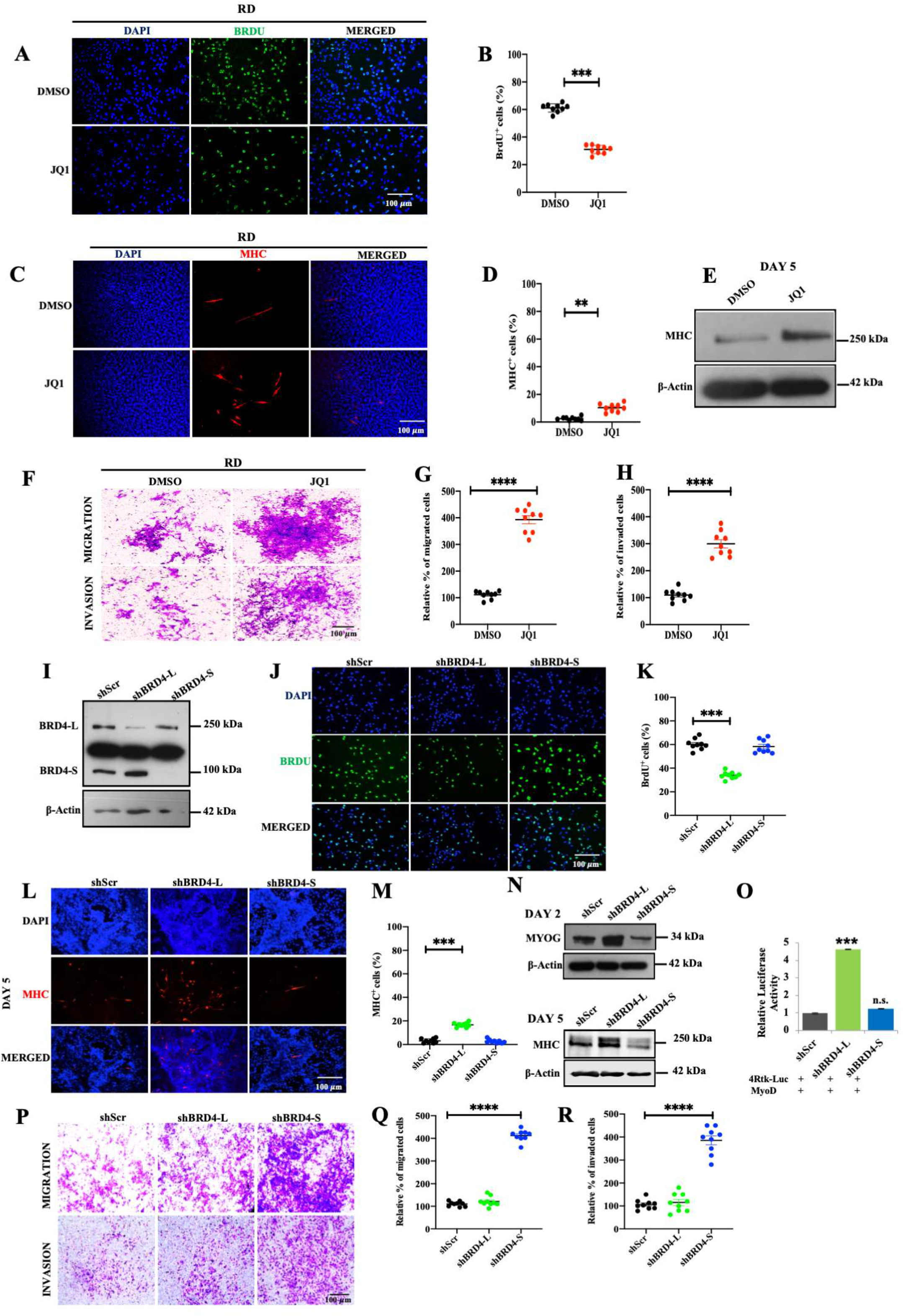
BRD4-L and BRD4-S have distinct roles in tumor growth and metastasis. **(A)** RD cells were treated with DMSO (vehicle) or 50 nM of JQ1 for 48 hr and proliferation was assessed using BrdU assay by immunofluorescence using anti-BrdU antibody. Nuclei were stained with DAPI (blue). Images are representative of at least three biological replicates. Scale bar: 100μm. **(B)** Scatter plot representing the percentage of BrdU^+^ cells in RD cells treated with DMSO or JQ1. Values correspond to the average ± SEM. Two-tailed non-parametric unpaired t test was performed for statistical analysis. ***p≤0.001. **(C)** RD cells were pretreated with 50 nM JQ1 in growth medium and then differentiated with DMSO or JQ1 for 5 days and analysed by immunofluorescence using anti-MHC antibody. Images are representative of at least three biological replicates. Scale bar: 100 µm **(D)** Scatter plot representing percentage of MHC^+^ cells in DMSO and JQ1-treated RD cells. The values correspond to average ± SEM. Two-tailed non-parametric unpaired t test was performed for statistical analysis. **p ≤ 0.01. **(E)** Western blot analysis using anti-MHC antibody in RD cells treated with DMSO or JQ1 in differentiation media for 5 days. β-Actin was used as loading control. **(F)** Migratory and invasive capacity of RD cells treated with 50 nM of JQ1 for 48 hr was assessed using transwell assays followed by crystal violet staining of the inserts. Images are representative of at least three biological replicates. Scale bar: 100 μm. **(G-H)** Scatter plot representing the percentage of migration and invasion of RD cells treated with DMSO or JQ1. The values correspond to average ± SEM. Two-tailed non-parametric unpaired t test was performed for statistical analysis. ****p ≤ 0.0001. **(I)** Stable knockdown of BRD4-L (shBRD4-L) and BRD4-S (shBRD4-S) in RD cells was analysed using Western blotting. β-Actin was used as loading control. Images are representative of at least three biological replicates**. (J)** Proliferation was assessed using BrdU assay in shScr, shBRD4-L and shBRD4-S RD cells by immunofluorescence using anti-BrdU antibody. Nuclei were stained with DAPI. Images are representative of at least three biological replicates. Scale bar: 100 μm. **(K)** Scatter plot representing the percentage of BrdU^+^ cells in shBRD4-L and shBRD4-S compared to shScr cells. The values correspond to average ± SEM. Two-tailed non-parametric unpaired t test was performed for statistical analysis. ***p ≤ 0.001. **(L)** shScr, shBRD4-L and shBRD4-S cells were cultured in differentiation media for 5 days and analysed by immunofluorescence using anti-MHC antibody. Nuclei were stained with DAPI. Images are representative of at least three biological replicates. Scale bar: 100 μm. **(M)** Scatter plot representing percentage of MHC^+^ cells in shBRD4-L and shBRD4-S cells compared to control cells. The values correspond to average ± SEM. Two-tailed non-parametric unpaired t test was performed for statistical analysis. ***p ≤ 0.001. **(N)** Western blot analysis of MYOG at Day 2, and MHC at Day 5 in shBRD4-L and shBRD4-S compared to shScr after culturing cells in differentiation media. β-Actin was used as loading control. **(O)** MyoD reporter activity was analysed in shScr, shBRD4-L and shBRD4-S cells by transfecting cells with 200 ng of the MRF reporter 4Rtk-luc, 200 ng MyoD and 5 ng Renilla. Luciferase activity was analyzed 48 hr post transfection (n=4). The values correspond to average ± SEM. Two-tailed non-parametric unpaired t test was performed for statistical analysis. ***p ≤ 0.001, n.s. not significant. **(P)** Migratory and invasive capacity of shScr, shBRD4-L and shBRD4-S cells was assessed using transwell assays followed by crystal violet staining of the inserts. Images are representative of at least three biological replicates. Scale bar: 100 μm. **(Q-R)** Scatter plot representing the percentage of migration and invasion of shScr, shBRD4-L and shBRD4-S cells. The values correspond to average ± SEM. Two-tailed non-parametric unpaired t test was performed for statistical analysis. ****p ≤ 0.0001. Numbers indicate the molecular weight of proteins.

To examine whether this effect was due to distinct roles of the two main BRD4 isoforms, BRD4-L and BRD4-S were individually silenced by stable knockdown in RD cells (**Fig. 2I**). The impact of isoform-specific depletion of BRD4 on proliferation, differentiation, migration and invasion was analysed. A significant decrease in the percentage of BrdU^+^ cells was apparent only in shBRD4-L cells compared to the control shScr cells. No significant change was observed in shBRD4-S cells (**Fig. 2J, K**). Similar results were observed with transient knockdown of the two isoforms in RD, RD18 and JR1 cell lines (Supplementary Fig. S1). A significant increase in Myosin heavy Chain (MHC)^+^ cells and of Myogenin (MYOG) and MHC expression was seen only in shBRD4-L cells compared to shScr, with no overt difference in shBRD4-S cells (**Fig. 2L-N**; Supplementary Fig. S2). Consistent with increased differentiation, shBRD4-L cell lines had a significantly high MRF reporter activity compared to shScr and shBRD4-S cells (**Fig. 2O**). Interestingly, a significant increase in migratory and invasive capacity was apparent only in shBRD4-S cells compared to shScr and shBRD4-L cells (**Fig. 2P, Q and R**; and Supplementary Fig. S3). These results demonstrate that BRD4-L promotes proliferation and inhibits myogenic differentiation, while BRD4-S inhibits migration and invasion of ERMS cells.

To validate these findings *in vivo*, RD control (shScr), shBRD4-L or shBRD4-S cells were injected subcutaneously in BALB/c nude mice. The tumour volume was similar between mice injected with shBRD4-S cells and shScr cells (**Fig. 3A and B**). On the other hand, a significant reduction in tumour volume was seen for the shBRD4-L group and tumors from only four mice could be resected. No adverse effect on the weight of the mice was noted in any group (**Fig. 3C**). We then tested the impact of the BRD4 isoforms in metastasis. Control shScr, shBRD4-L and shBRD4-S cells were injected into NOD/SCID mice through the tail vein. The control and shBRD4-L group showed a similar number of tumors in the liver. On the other hand, mice injected with shBRD4-S cells showed a strikingly higher number and size of nodules primarily in the liver but also in the kidney, stomach and lungs (**Fig. 3D and E).** No apparent difference was observed in the body weight of any group (**Fig. 3F**). The liver, lung and kidney nodules were analyzed histologically. Widespread metastasis in the liver, lungs and kidney was apparent in mice injected with shBRD4-S cells compared to control and shBRD4-L group (Supplementary Fig. S4).

**Figure 3:**
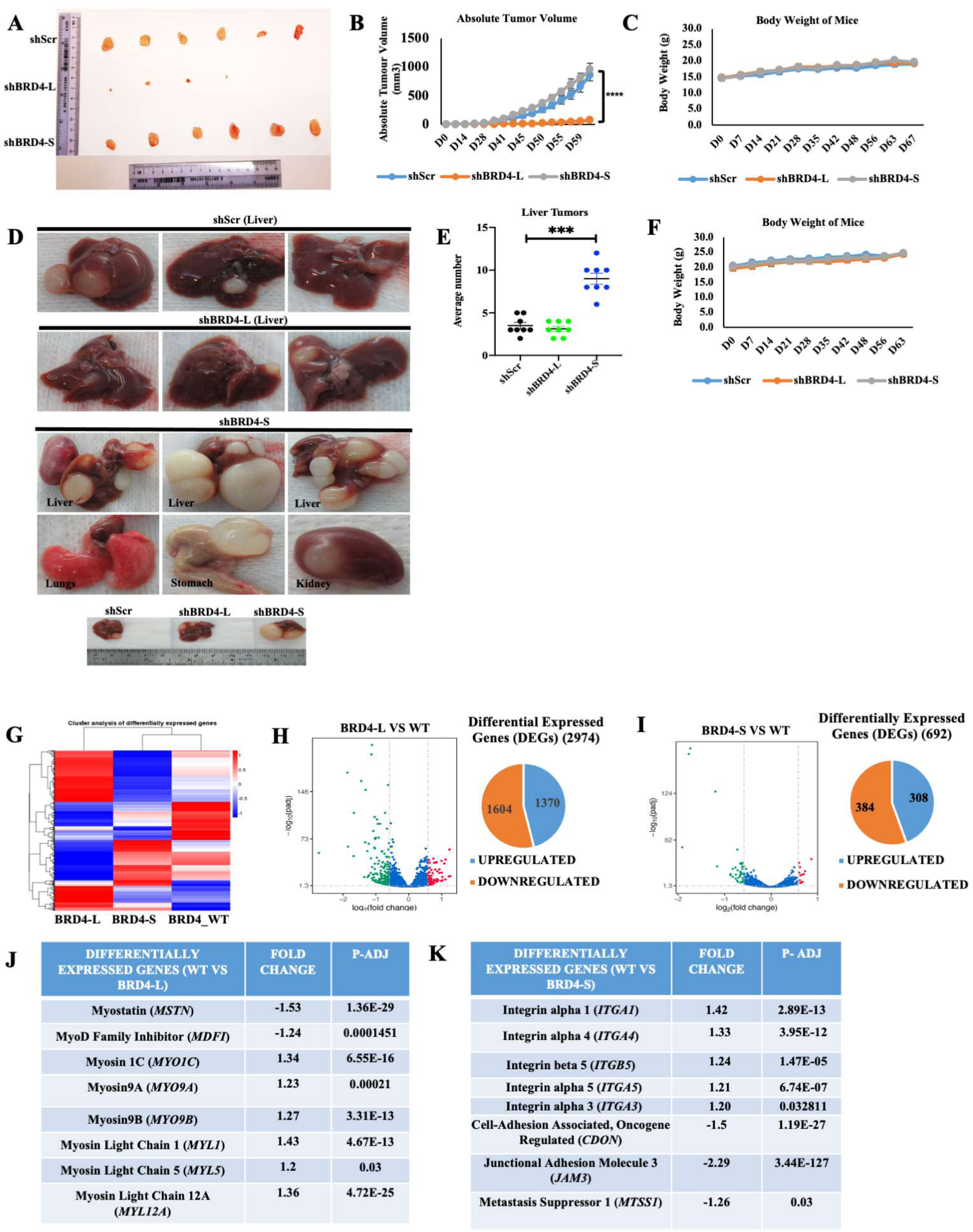
BRD4-L enhances tumor growth whereas BRD4-S blocks metastasis *in vivo.* **(A)** Nude mice (n=10/group) were injected with shScr, shBRD4-L and shBRD4-S cells to analyse tumor growth. Tumors were resected from 6 shScr and shBRD4-S mice and from 4 shBRD4-L mice. **(B-C)** The graphs represent absolute tumor volume **(B)** and average body weight **(C)** of mice. Statistical significance was calculated using repeated-measure one-way ANOVA where ****p ≤ 0.0001. Values correspond to the average ± SEM. **(D)** NOD/SCID mice were injected through the tail vein with shScr, shBRD4-L and shBRD4-S cells (n=10/group). Metastasis was seen in the liver for shScr and shBRD4-L groups, while tumors were seen in the liver, lungs, stomach and kidney in shBRD4-S group. **(E)** Scatter plot representing the average number of liver nodules in mice injected with shScr, shBRD4-L and shBRD4-S cells (n=8). Values correspond to the average ± SEM. Two-tailed non-parametric unpaired t test was performed for statistical analysis. ***p ≤ 0.001. **(F)** Graph representing average body weights of control shScr, shBRD4-L, shBRD4-S mice. Values correspond to the average ± SEM. **(G)** Heat map indicates hierarchical cluster of differentially expressed genes (DEGs) between control siScr (BRD4_WT), siBRD4-L and siBRD4-S RD cells. Red and blue colours represents high and low expression of genes respectively. (**H-I**) Volcano plot and pie charts indicate the distribution and number of upregulated and downregulated genes in siBRD4-L versus control (WT) and siBRD4-S compared to siScr control (WT) groups. (**J**) A list of the top significantly altered genes related to myogenesis in the siBRD4-L vs siScr group is shown with the fold change and *p*-adjusted values. (**K**) A list of the top significantly altered genes related to migration and invasion identified in the siBRD4-S vs siScr group with the fold change and *p*-adjusted values.

### BRD4-L and BRD4-S regulate distinct target genes

To identify the mechanisms underlying their distinct roles, we performed RNA-sequencing analysis using three technical replicates of siScr, siBRD4-L and siBRD4-S RD cells. Hierarchical cluster analysis of differentially expressed genes (DEGs) and volcano plots showed that 2974 and 692 genes were differentially regulated in siBRD4-L and siBRD4-S cells respectively when compared to BRD4_WT (**Fig. 3G, H and I**). 1370 genes were upregulated while 1604 genes were downregulated in siBRD4-L cells (**Fig. 3H**). 308 genes were upregulated and 384 genes were downregulated in siBRD4-S cells (**Fig. 3I**). Gene Ontology (GO) analysis showed that cell-substrate adherens junction, focal adhesion and actin cytoskeleton were upregulated in siBRD4-L cells that are important for cellular differentiation. Similarly, cell-substrate adherens junction, and focal adhesion were upregulated with siBRD4-S cells correlating with cell motility (Supplementary Fig. S5A). In corroboration with the phenotypic effects, myogenic differentiation genes were modulated upon BRD4-L knockdown. Several genes, including *MSTN, MDFI,* were downregulated while *MYO1C, MYO9A, MYO9B, MYL1, MYL5 and MYL12A* were upregulated on silencing BRD4-L (**Fig. 3J**). In contrast, upon BRD4-S knockdown, a distinct set of genes that promote migration and invasion including integrins *ITGA1* (*29*)*, ITGA3* (*30*)*, ITGA4* (*31*)*, ITGA5* (*32*) were upregulated while some metastasis-suppressors like *CDON, MTSS1 and JAM3* were downregulated (**Fig. 3K**). We validated some of the targets regulated by BRD4-L and BRD4-S by RT-qPCR. The mRNA level of *MSTN* was downregulated while *MYL12A* and *MYL5* were upregulated in siBRD4-L cells compared to siScr and siBRD4-S cells (Supplementary Fig. S5B). On the other hand, expression of *ITGA1*, *ITGA3*, *ITGA4* and *ITGA5* was repressed in siBRD4-S cells compared to siScr and siBRD4-L cells (Supplementary Fig. S5C).

### BRD4-L regulates myogenic differentiation through myostatin

Among the genes that were downregulated upon BRD4-L knockdown, we were particularly interested in myostatin (*MSTN*) that is a negative regulator of myogenic differentiation in RMS cells (33). Myostatin protein was downregulated in shBRD4-L cells compared to shScr and shBRD4-S cells (**Fig. 4A**). Moreover, tumour lysates from two independent mice from the xenograft assays showed that MHC levels were elevated and myostatin levels were significantly reduced in shBRD4-L tumours compared to shScr control and shBRD4-S tumours (**Fig. 4B**). To examine whether BRD4-L directly regulates myostatin, we examined BRD4-L occupancy at the myostatin promoter using ChIP-qPCR. A significant occupancy of BRD4-L was seen at the myostatin promoter with no enrichment of BRD4-S (**Fig. 4C**). Moreover, enrichment of the activation mark H3K9Ac (**Fig. 4D**) along with RNA Pol II (Ser2P) (**Fig. 4E**) was seen. No significant change in enrichment of BRD4-L and RNA Pol II was seen in shBRD4-S cells compared to control cells (**Fig. 4F and G**).

**Figure 4:**
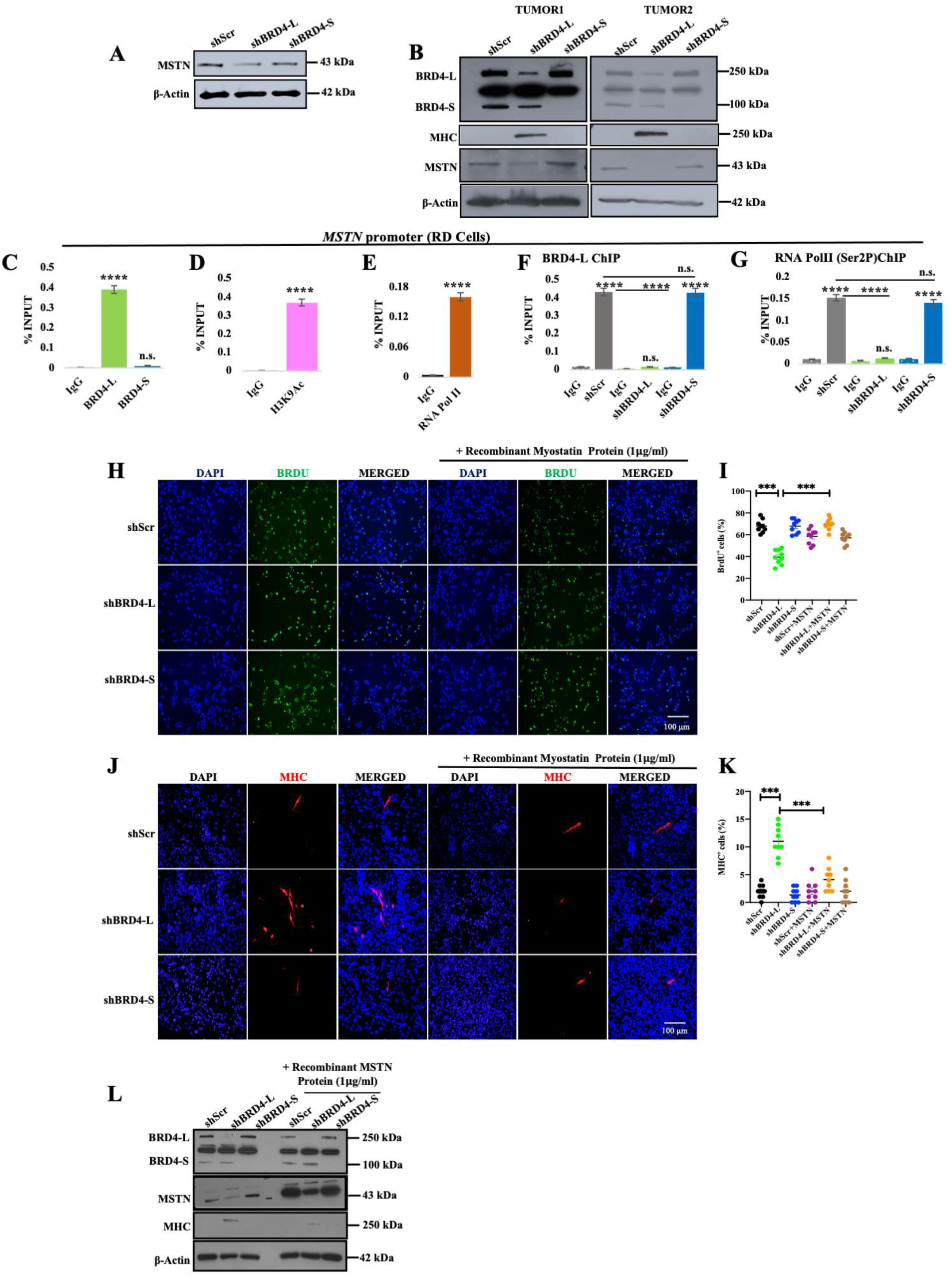
BRD4-L promotes proliferation and represses myogenic differentiation via regulation of MSTN expression. **(A)** MSTN protein level was analyzed in shScr, shBRD4-L and shBRD4-S cells. β-Actin was used as loading control. Images are representative of at least three biological replicates. **(B)** Tumor lysates from 2 independent RD shScr, shBRD4-L and shBRD4-S xenografts were analysed for BRD4, MHC and MSTN expression using Western blotting. Β-Actin was used as loading control. **(C-E)** ChIP assay was performed to examine the enrichment of BRD4-L and BRD4-S isoforms **(C)**, H3K9Ac **(D)** and RNA Pol II (Ser2P) **(E)** on the *MSTN* promoter relative to IgG control (n=3). The values correspond to the average ± SEM. Two-tailed non-parametric unpaired t test was performed for statistical analysis**. (F-G)** ChIP assay to examine the enrichment of BRD4-L **(F)** and RNA Pol II (Ser2P) **(G)** in shBRD4-L and shBRD4-S compared to shScr control cells on MSTN promoter (n=4). The values correspond to average ± SEM. Two-tailed non-parametric unpaired t test was performed for statistical analysis. ****p ≤ 0.0001, n.s. not significant. **(H)** BrdU assay in shScr, shBRD4-L and shBRD4-S cells and treated with or without 1 μg/ml of recombinant myostatin protein for 72 hr. Nuclei were stained with DAPI (blue). Images are representative of at least three biological replicates. Scale bar: 100 μm. **(I)** Scatter plot showing the percentage of BrdU^+^ cells in shBRD4-L, shBRD4-S in comparison with shScr cells treated with or without recombinant myostatin protein. The values correspond to the average ± SEM. Statistical significance was calculated by one-way ANOVA analysis. ***p ≤ 0.001. **(J)** shScr, shBRD4-L and shBRD4-S cells were cultured in differentiation media in the absence or presence of 1μg/ml of recombinant myostatin protein for 72 hr. MHC^+^ cells were analysed by immunofluorescence with anti-MHC antibody. Images are representative of at least three biological replicates. Scale bar: 100 μm. **(K)** Scatter plot indicating MHC^+^ cells in shBRD4-L, shBRD4-S in comparison with shScr cells with or without recombinant myostatin protein. Statistical significance was calculated by one-way ANOVA analysis. ***p ≤ 0.001. **(L)** Western blot analysis indicating protein expression of MHC and MSTN in shScr, shBRD4-L and shBRD4-S cells with or without recombinant myostatin protein for 72 hr. β-Actin was used as loading control.

To confirm that myostatin is a downstream effector of BRD4-L, we added exogenous recombinant myostatin protein to shBRD4-L cells. Addition of myostatin increased proliferation in shBRD4-L cells (**Fig. 4H and I**). A modest decrease in proliferation in shScr and shBRD4-S cells was apparent on addition of myostatin. Myostatin also reversed the enhanced myogenic differentiation seen in shBRD4-L cells (**Fig. 4J and K**) that was evident by the decrease in MHC^+^ cells **(Fig. 4K)** and by Western blot analysis (**Fig. 4L**).

### An interplay between BRD4-S and BRD4-L is involved in regulation of integrin genes

Several integrin genes (*ITGA1, ITGA3, ITGA4, ITGA5, ITGB5*) were significantly upregulated upon knockdown of BRD4-S (**Fig. 3K and** Supplementary Fig. S5B). We validated ITGA4 and ITGA5 protein expression in shBRD4-S cells compared to shBRD4-L and shScr control cells (**Fig. 5A**). Additionally, a remarkably high expression of ITGA4 and ITGA5 was seen in the liver nodules isolated from two independent shBRD4-S mice from the *in vivo* metastasis experiments (**Fig. 5B**). Since BRD4 mostly functions as an activator, we examined how loss of BRD4-S could result in an upregulation of integrin genes. A higher enrichment of BRD4-S compared to BRD4-L was apparent at *ITGA1, ITGA3, ITGA4, ITGA5* genes (**Fig. 5C**). A significant enrichment of H3K9Ac was also seen at the promoters of *ITGA4* and *ITGA5* (**Fig. 5D**) although occupancy of RNA Pol II was not observed (**Fig. 5E**).

**Figure 5:**
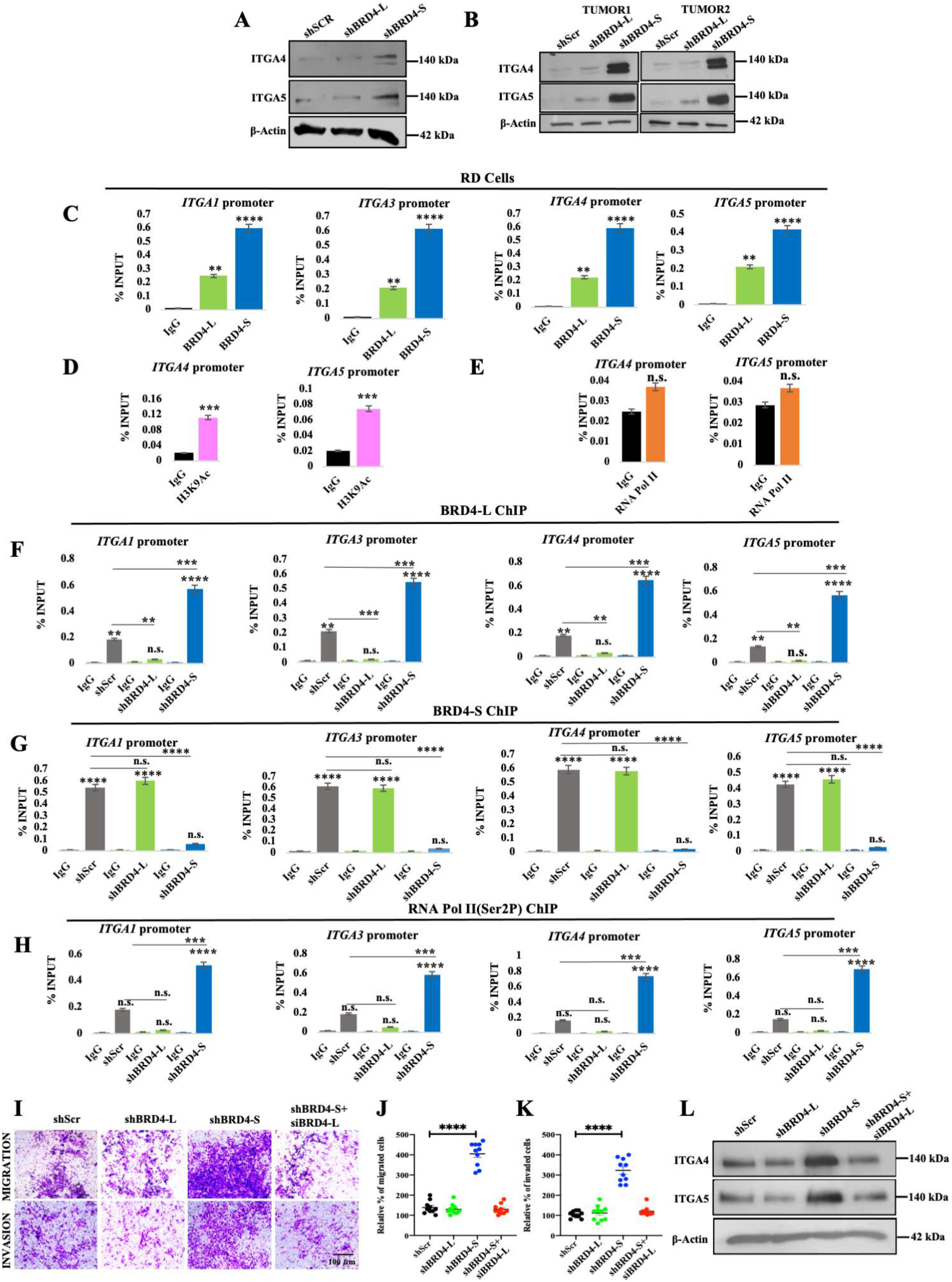
Integrins are downstream targets of BRD4-S. **(A)** Western blot analysis of ITGA4 and ITGA5 in shScr, shBRD4-L and shBRD4-S cells. β-Actin was used as loading control. Images are representative of at least three biological replicates. **(B)** Tumor lysates from 2 independent NOD/SCID mice from tail vein assays were analysed for expression of ITGA4 and ITGA5 using western blotting. β-Actin was used as loading control. **(C)** ChIP assay was performed to examine the enrichment of BRD4-L and BRD4-S on *ITGA1*, ITGA3, *ITGA4* and *ITGA5* promoter, IgG was used as a control (n=3). The values correspond to the average ± SEM. Two-tailed non-parametric unpaired t test was performed for statistical analysis. **p ≤ 0.01, ****p ≤ 0.0001. **(D-E)** ChIP assay was performed to examine the enrichment of H3K9Ac **(D)** and RNA Pol II (Ser2P) **(E)** on the *ITGA4* and *ITGA5* promoter (n=3). The values correspond to average ± SEM. Two-tailed non-parametric unpaired t test was performed for statistical analysis. ***p ≤ 0.001, n.s. = not significant. **(F-H)** Enrichment of BRD4-L **(F)**, BRD4-S **(G)** and RNA Pol II (Ser2P) **(H)** in shScr, shBRD4-L and shBRD4-S cells at the *ITGA1*, *ITGA3*, *ITGA4* and *ITGA5* promoter was analyzed by ChIP assay (n=4). The values correspond to average ± SEM. Two-tailed non-parametric unpaired t test was performed for statistical analysis. n.s. not significant, **p ≤ 0.01, ***p ≤ 0.001 and ****p ≤ 0.0001. **(I)** Upon transient knockdown of BRD4-L in shBRD4-S cells for 48 hr, migratory and invasive capacity of shScr, shBRD4-L and shBRD4-S cells was analyzed using transwell assay. The inserts were stained with crystal violet. Images are representative of at least three biological replicates. Scale bar: 100 μm. **(J-K)** Scatter plots shows the percentage of migrated **(J)** and invaded **(K)** shScr, shBRD4-L, shBRD4-S and shBRD4-S + siBRD4-L cells after 48 hr. Statistical significance was calculated by one-way ANOVA analysis. ****p ≤ 0.0001. **(L)** Western blot analysis of ITGA4 and ITGA5 in shScr, shBRD4-L, shBRD4-S and shBRD4-S + siBRD4-L. β-Actin was used as loading control.

Previous studies have demonstrated that a BRD4-S-like activity via deletion of the proline-rich region in BRD4-L may oppose the function of BRD4-L by competing for target acetyl lysine residues (34). We therefore examined if BRD4-S inhibits binding of BRD4-L at the integrin gene promoters. Interestingly, in the shBRD4-S cells, an increase in BRD4-L enrichment was seen, indicating that reduction in BRD4-S allows for elevated BRD4-L occupancy (**Fig. 5F**). Notably, there was no significant change in BRD4-S enrichment in shBRD4-L cells compared to shScr control cells (**Fig. 5G**). In addition to elevated BRD4-L occupancy, there was also enrichment of RNA Pol II in shBRD4-S cells, indicating an assembly of a transcription complex (**Fig. 5H**).

To confirm whether BRD4-L drives expression of integrin genes and cellular motility in the absence of BRD4-S, we silenced BRD4-L using siRNA in the shBRD4-S cells. Interestingly, the migratory and invasive capacity of the shBRD4-S plus siBRD4-L cells was reduced to that of shScr cells (**Fig. 5I, J and K**). Western blot analysis revealed that expression of ITGA4 and ITGA5 in shBRD4-S cells transfected with siBRD4-L was similar to that of shScr control cells (**Fig. 5L**). To confirm whether ITGA4 and ITGA5 are key targets involved in increased migration and invasion of shBRD4-L cells, we transiently knocked down their expression individually or together in shScr, shBRD4-L and shBRD4-S cells. Depletion of ITGA4 or ITGA5 individually partially reversed the phenotype of shBRD4-S cells, but the reduction was very pronounced when both genes were silenced in shBRD4-S cells (Supplementary Fig. S6A-E).

Collectively, our data support a model where BRD4-L is involved in tumor progression and inhibits differentiation of ERMS cells, at least in part, by regulation of myostatin. BRD4-S, on the other hand, acts as a gatekeeper. Removal of the blockade imposed by BRD4-S permits BRD4-L to promote metastasis by regulating expression of integrin genes.

## Discussion

In this study, we provide the first evidence for distinct isoform-specific roles of BRD4 in ERMS. We demonstrate that BRD4-L represses MyoD activity and myogenic differentiation. On the other hand, intriguingly BRD4-S limits the oncogenic role of BRD4-L by acting as a gatekeeper of metastasis. In the absence of BRD4-S, significant enrichment of BRD4-L as well as RNA Pol II is apparent on integrin gene promoters, resulting in their elevated expression that correlates with increased metastasis.

BRD4 is deregulated in various cancers including NUT Midline Carcinoma (NMC), Acute Myeloid Leukemia (AML), pancreatic, myeloma, lung and triple-negative breast cancer (TNBC) where it promotes oncogenesis (8,35). Consequently, pan-BET inhibitors (BETi) have been tested and show promising anti-tumour activity in diverse clinical models albeit with some limitations (16,19,36). For instance, while I-BET151 is effective at inhibiting primary tumour growth, it does not reduce pulmonary metastasis in orthotopic models of metastasis (37). Sensitivity to BETi has been suggested to largely, though not exclusively, depend on the amplification of MYC family members in multiple tumours (38–40). Conversely, BRD4 also functions as a tumour suppressor in some contexts. Its overexpression *in vivo*, caused a drastic reduction in tumor growth along with pulmonary metastasis (41). Similarly, BRD4 mediates resistance to transformation in patients with Hutchinson Gilford Progeria Syndrome (42). BET protein inhibitors such as JQ1 have shown promising results for the treatment of different cancers. However, JQ1 can have an undesired effect of promoting cancer metastasis (20,43) although it inhibited cancer cell growth (44). Similar to these studies, in ERMS cells, JQ1 reduced proliferation and increased differentiation but promoted migration and invasion.

Recent studies revealed opposing roles of the two isoforms, where BRD4-S functions as an oncogene, while BRD4-L acts as a tumour suppressor in breast cancer (12). Although BRD4-L functions as tumour and metastasis suppressor in breast cancer, unexpectedly, the P-TEFb-binding domain of BRD4-L, upon deletion of the proline-rich region, may enhance metastasis and induces EMT and cancer stem cell-like properties (34). Unlike breast cancer, where BRD4-L functions as a tumor suppressor, in ERMS, it appears to have an oncogenic role. These observations suggest that the two BRD4 isoforms may have opposing roles in different contexts that have not been systematically analyzed in most cancers.

Our data demonstrate that BRD4-L regulates differentiation at least in part through regulation of myostatin. Myostatin is a well-established negative regulator of myogenesis and inhibitor of MyoD activity. Myostatin has previously been shown to inhibit proliferation of RMS cells, and its suppression with a dominant negative form of activin receptor type IIb promotes differentiation. BRD4 interacts with the methyltransferase SMYD3 to regulate myostatin expression (45). On the other hand, our functional and transcriptomic data demonstrate that BRD4-S loss enhances expression of integrins due to increased BRD4-L binding to integrin gene promoters. These findings are consistent with previous studies which demonstrated that a BRD4-S-mimicking activity may compete with BRD4-L for binding to acetylated histones and interferes with BRD4-L function (34) likely due to an expanded histone binding of BRD4-S versus BRD4-L.

Our finding that each BRD4 isoform regulates specific gene targets is in line with previous studies (12,13). We identified several integrins which are key players in cellular adhesion and migration (46) among the targets that are regulated by BRD4-S. Integrins participate in the remodeling of ECM and colonization of cancer cells in new metastatic sites. N-cadherin and α9-integrin link the Notch pathway to cell adhesion, motility and invasion in RMS (47). The BRD4/c-Myc axis also intersects with integrin/FAK-dependent pathway in TNBC (48).

The mechanisms by which the two isoforms regulate specific gene transcription needs further investigation. Isoform-specific binding partners might determine binding specificity and distinct functions of the isoforms. Taken together, our study highlights the value of BRD4-isoform specific therapeutic strategies and indicates that BRD4-S expression may be a biomarker of cancer metastasis.

## Data Availability

The RNA-Seq data has been deposited in GEO under the accession number GSE215393. The private token for reviewers is gtwnayygptkrjwr.

## Acknowledgements

We thank Peter Houghton and Rosella Rota for the ERMS cell lines. This work was supported by a Ministry of Education MOE2019-T2-1-024 grant to R. Taneja. C.-M. Chiang’s research is supported by US NIH grant 1RO1CA251698-01 and CPRIT grant RP180349. Jia Yu Leung was supported by Agency for Science, Technology and Research(A*STAR). We thank Dr. Nandini Karthik and Dr. Hsin Yao Chiu for comments on the manuscript.

## Authors’ Contributions

**Conception and design:** D Das and R Taneja

**Development of methodology:** D Das

**Acquisition of data (provided animals, acquired and managed patients, provided facilities, etc.):** D Das, JY Leung and AHP Loh

**Analysis and interpretation of data (e.g., statistical analysis, biostatistics, computational analysis):** D Das, APH Loh, V Tergaonkar and R Taneja

**Writing, review, and/or revision of the manuscript:** D Das R Taneja and C-M Chiang

**Study supervision:** R Taneja

